# Indigenous generalist pollinators shape plant–insect interaction networks across a land-use gradient on an oceanic island

**DOI:** 10.64898/2026.10.17.759801

**Authors:** François Rigal, Ana Picanço, Sérgio Timóteo, Mário Boieiro, Paulo A. V. Borges

**Affiliations:** Université de Pau et des Pays de l’Adour, CNRS, Institut des Sciences Analytiques et de Physico-Chimie pour l’Environnement et les Matériaux (IPREM), Pau, France; University of the Azores, Centre for Ecology, Evolution and Environmental Changes (CE3C) & CHANGE—Global Change and Sustainability Institute, Rua Capitão João d’Ávila, Pico da Urze, 9700-042 Angra do Heroísmo, Portugal; Centre for Functional Ecology, Associate Laboratory TERRA, Department of Life Sciences, University of Coimbra, Calçada Martim de Freitas, Coimbra 3000-456, Portugal; Centre for Ecology, Evolution and Environmental Changes (CE3C) & CHANGE—Global Change and Sustainability Institute, Faculty of Sciences, University of Lisbon, Lisbon, Portugal; LIBRe—Laboratory for Integrative Biodiversity Research, Finnish Museum of Natural History, University of Helsinki, 00014 Helsinki, Finland; IUCN SSC Atlantic Islands Invertebrate Specialist Group, 9700-042 Angra do Heroísmo, Azores, Portugal; IUCN SSC Species Monitoring Specialist Group, 9700-042 Angra do Heroísmo, Azores, Portugal

**Keywords:** Indigenous species, introduced species, flower-visiting insects, land-use change, nestedness, modularity, oceanic islands, pollination networks, species role

## Abstract

1. The study asks how plant–pollinator networks reorganise along a land-use gradient spanning five habitat types, ranging from native forests to intensively managed pastures, on the oceanic island of Terceira (Azores, Portugal).
2. Interaction dissimilarity between habitats was always very high (on average 87%) and this reshuffling was driven mainly by plant species turnover rather than insect turnover or partner rewiring.
3. Plant composition shifted strongly along the gradient, with introduced species increasingly dominating disturbed habitats, whereas pollinator richness and identity remained comparatively stable, consistent with more homogeneous communities dominated by widespread generalist indigenous insect species.
4. Connectance and nestedness were lowest, while modularity was highest, in intermediate habitats (naturalized vegetation, exotic forest) a pattern interpreted as the signature of mixed native and introduced floristic pools generating distinct modules, reinforced by locally selective insect foraging.
5. By identifying the species role in the landscape, nearly half of the flower-visiting insects was found to act as habitat connectors linking networks across the gradient. These connectors were mostly indigenous species that behaved as context-dependent generalists, switching plant partners between habitats rather than tracking the same resource everywhere.
6. Widespread pollinators with broad feeding spectra mediated the integration of introduced plants into local pollination networks. This highlights the key role of indigenous super-generalist pollinators in maintaining network resilience across the disturbance gradient. Nevertheless, the reliance on a few super-generalist species for pollination across diverse habitat-types, as reported from several archipelagos worldwide, pinpoints the vulnerability of these ecosystems to pollinator declines.

## INTRODUCTION

Biodiversity is declining rapidly due to various anthropogenic pressures, posing serious threats to the complex networks of ecological interactions that support community structure and function (Davison et al., 2021; Vilà & Hulme, 2017). Although human activities affect ecosystems worldwide, oceanic islands-those never connected to continental landmasses, are especially vulnerable (Fernández-Palacios et al. 2021). Their biotas, often simplified and evolutionarily naïve due to long-term geographic isolation, are especially susceptible to human disturbances, making these islands global hotspots for both species extinctions and biological invasions (Arjona et al., 2025; Brooks et al., 2002; Kueffer et al., 2010; Matthews et al., 2024). In this context, plant–pollinator networks and other mutualistic interactions on oceanic islands have emerged over the past decades as key research areas, from both conservation and restoration perspectives (Heinen et al., 2023; Kaiser-Bunbury et al., 2010; Traveset et al., 2013), and as model systems for understanding how anthropogenic disturbance can reshape the structure, dynamics, and stability of ecological networks (Trøjelsgaard et al., 2019; Vizentin-Bugoni et al., 2019).

Few consistent patterns have emerged from studies of pollination networks on oceanic islands. Compared with mainland networks, island networks are generally smaller and simpler, with lower interaction diversity and the absence of certain functional groups, such as long-tongued pollinators and social bees (Castro-Urgal & Traveset, 2014; Traveset et al., 2013; Trøjelsgaard & Olesen, 2013). A striking feature of island networks is the presence of indigenous super-generalist species (Boieiro et al., 2025a; Hayes et al., 2019; Kaiser-Bunbury et al., 2009; Olesen et al., 2002). This concept was first introduced by Olesen et al. (2002), based on observations from Flores (Azores) and Île aux Aigrettes (Mauritius), where endemic pollinator insects showed exceptionally broad pollination niches. Similar patterns have since been reported across other archipelagos (e.g. Crichton et al., 2019). In Fiji, for instance, the native bee *Homalictus fijiensis* exploits a wider range of floral hosts than introduced species such as *Apis mellifera* (Draper et al., 2021). Super-generalism is often attributed to ecological release and density compensation (Olesen et al., 2002), and is thought to shape key structural properties of island networks, including strong interaction asymmetry, high connectance, high nestedness, and low modularity (Dupont et al., 2024; Olesen & Jordano, 2002; Olesen et al., 2007; Traveset et al. 2015; Trøjelsgaard & Olesen, 2016).

The ecological context of island pollination networks has been profoundly transformed due to human-driven changes, particularly in the last few centuries (Fernández-Palacios et al. 2021). Many oceanic islands have undergone extensive habitat loss, with native ecosystems replaced by mosaics of anthropogenic land uses differing in disturbance intensity (Borges et al., 2018; Caujapé-Castells et al., 2010). These land-use changes go hand in hand with the introduction of non-indigenous species (Kremen et al., 2007; Leclerc et al., 2018; Traveset & Richardson, 2014; Tylianakis et al., 2008), leading to the emergence of novel pollination networks where indigenous and introduced species coexist and interact to form new ecological assemblages (Traveset et al., 2013; Trøjelsgaard et al., 2019). Yet, how these novel networks differ from those in relatively undisturbed habitats and how indigenous super-generalist pollinators establish their roles within these newly formed assemblages remain poorly understood (Burkle & Alarcón, 2011; Kaiser-Bunbury & Blüthgen, 2015). In this context, indigenous super-generalist pollinators may play a pivotal role in mediating interactions across different land-use types. On the one hand, such species may facilitate the persistence and spread of introduced plants across habitats (Traveset & Richardson, 2014); on the other, they might enhance the survival of indigenous plant populations that persist within and outside their original habitat (Ghisbain et al., 2021). Moreover, by maintaining interaction links among diverse species and land-use types, indigenous super-generalist pollinators could substantially contribute to the structural stability, resilience, and functional continuity of pollination networks in anthropogenic landscapes (Memmott et al., 2004).

In this study, we examined plant–pollinator networks across five distinct land-use types representing a gradient of management intensity on Terceira Island (Azores, Portugal). The Azores, and particularly Terceira, provide a good model system for investigating the structure and dynamics of pollination networks on oceanic islands. The archipelago harbours one of the best-documented floras and faunas among island systems, encompassing both indigenous and introduced species (Borges et al., 2010; Borges et al. 2022; Leo et al., 2024; Pozsgai et al., 2024). Today, Azorean islands are covered by a mosaic of land-use types with varying levels of disturbance, ranging from native forests to intensively managed pastures (Gaspar et al., 2008; Rego et al., 2015; Triantis et al., 2010). Recent studies on Terceira Island have highlighted the presence of several pollinator species displaying broad habitat distributions and generalized feeding habits, thus supporting the presence of super-generalist dynamics within the island networks (Boieiro et al., 2025b; Picanço et al., 2017a; Weissmann & Schaefer, 2017).

Herein, we investigate how plant–pollinator networks respond to land-use change on Terceira Island (Azores) by (1) examining how interaction patterns and network structure vary along the land-use gradient, and (2) identifying the structural roles of pollinator insect and plant species across this gradient, with particular attention to the potential role of indigenous super-generalist pollinators. We hypothesize a strong turnover in species and interaction composition among networks from different land-use types, particularly between remnant native forest patches and intensively-managed land-uses. Based on previous studies in the Azores, plant communities change composition between land-uses (Picanço et al., 2017a), with native species being gradually replaced by introduced ones as disturbance intensity increases. In contrast, pollinator communities are likely to persist across habitats due the presence of super-generalists that will shift their interaction partners with changing plant communities. Consequently, as the dominance of super-generalist pollinators increases and specialist species decrease along the land-use gradient, we expect a decline in overall network specialization, an increase of nestedness and a decrease of modularity (Proesmans et al., 2024).

## MATERIALS AND METHODS

### Study area

The Azores (37° to 40° N; 25° to 31° W) are one of the world’s most isolated archipelagos, comprises nine main volcanic islands aligned on a WNW–ESE axis in the Atlantic Ocean. Before human colonization, around AD 1440, the Azorean archipelago was almost entirely forest-covered (Elias et al., 2016). By c. AD 1850, human activities had restricted the native forest to areas above 500 m a.s.l., while during the 20^th^ century, the development of a dairy-based economy has led to a further reduction of the native forest, with the clearing of large fragments at mid- and high-elevations for pasture development. In addition, the plantation of forests of exotic species was incentivized during the 1950s in areas unsuitable for cattle grazing, thus promoting additional deforestation of natural forest habitat. Thus, in < 600 years, 95% of the original native forest has been destroyed (Gaspar et al., 2008).

Our study was conducted on Terceira, a relatively young island (3.52 Ma) with 402 km^2^ of surface area, where the native forest is now restricted to 23 km^2^, about 6% of its original area (Triantis et al., 2010 and references herein). The non-urban landscape of Terceira is dominated by five major habitats: (i) native forests dominated by the endemic *Erica azorica*, *Juniperus brevifolia* and *Laurus azorica*, (ii) naturalized vegetation combining native and exotic flora (*Calluna vulgaris*, *Erica azorica*, *Hedychium gardnerianum* and *Pittosporum undulatum)* (iii) exotic forests (*Cryptomeria japonica* and *Eucalyptus globulus* monoculture plantations, monocultures and mixed forest of the invasive *Pittosporum undulatum*); (iv) semi-natural pastures (mid- and high-elevation pastures maintaining some indigenous plants) subject to short-term cattle grazing activity and low inputs of fertilizers and (v) intensively managed pastures subject to intensive cattle grazing and high inputs of fertilizers. The five land-uses were ranked in order of increasing land-use intensity, with (1) native forest, (2) naturalized vegetation, (3) exotic forest, (4) semi-natural pasture and (5) intensively managed pasture (see Cardoso et al., 2013; Picanço et al., 2017a; Rigal et al., 2018 and Table S1 for detailed features of the land-uses).

### Sampling methodology

Field work was conducted from June to September 2013 and from July to October 2014, a period when most plant species are flowering and with the most favourable weather conditions for the activity of flower-visiting insects. Fifty sites were selected (10 sites per habitat type) and, in each, a 10 m long linear transect (1 m width) was set up (Figure 1A). To select the 10 sites per habitat type, we maximize the covered environmental diversity following Jiménez-Valverde and Lobo (2004) and Aranda et al. (2011). First, an environmental matrix for Terceira Island was compiled using climatic, topographic and geological variables with a resolution of 100 x 100 m. Using the *k*-means non-hierarchical clustering algorithm, we grouped all cells of each habitat type in 10 clusters, making a total of 50 clusters (5 habitats x 10 groups). For each cluster, we ordered the cells according to their distance to the group’s multidimensional centroid using Euclidean distance. The first cell in this ranking, deemed to be the most representative of the cluster, was chosen for sampling. If it was impossible to reach the selected cell in the field due to inaccessibility or lack of authorization from land owners, the second cell was chosen and so forth. Transect surveys were carried out once a year in both 2013 and 2014, and the order of surveys to the transects were randomized. Each survey took place during sunny days between 09:00 and 18:00 and lasted 180 minutes. Within each site, transects were positioned purposively to encompass actively flowering patches and thereby obtain sufficient observations of plant– flower-visitor interactions. This design targeted interaction structure rather than habitat-wide abundance, and sampling effort and transect dimensions were standardised across sites. Along the transect, randomly selected flowers were surveyed for 4 minutes each, and all insects probing for nectar or eating/collecting pollen were recorded. However, we acknowledge that the timed point sampling method may overestimate fast-moving visitor species and those exhibiting plant fidelity. Flower-visiting insects were collected with an entomological net to a pooter, if it was not possible to identify them in the field. Note that here we use the term ‘pollinator’ regardless of its effectiveness in the pollination process, as we do not know whether flower visitation results in pollination (Traveset et al. 2015).

**Figure 1.**
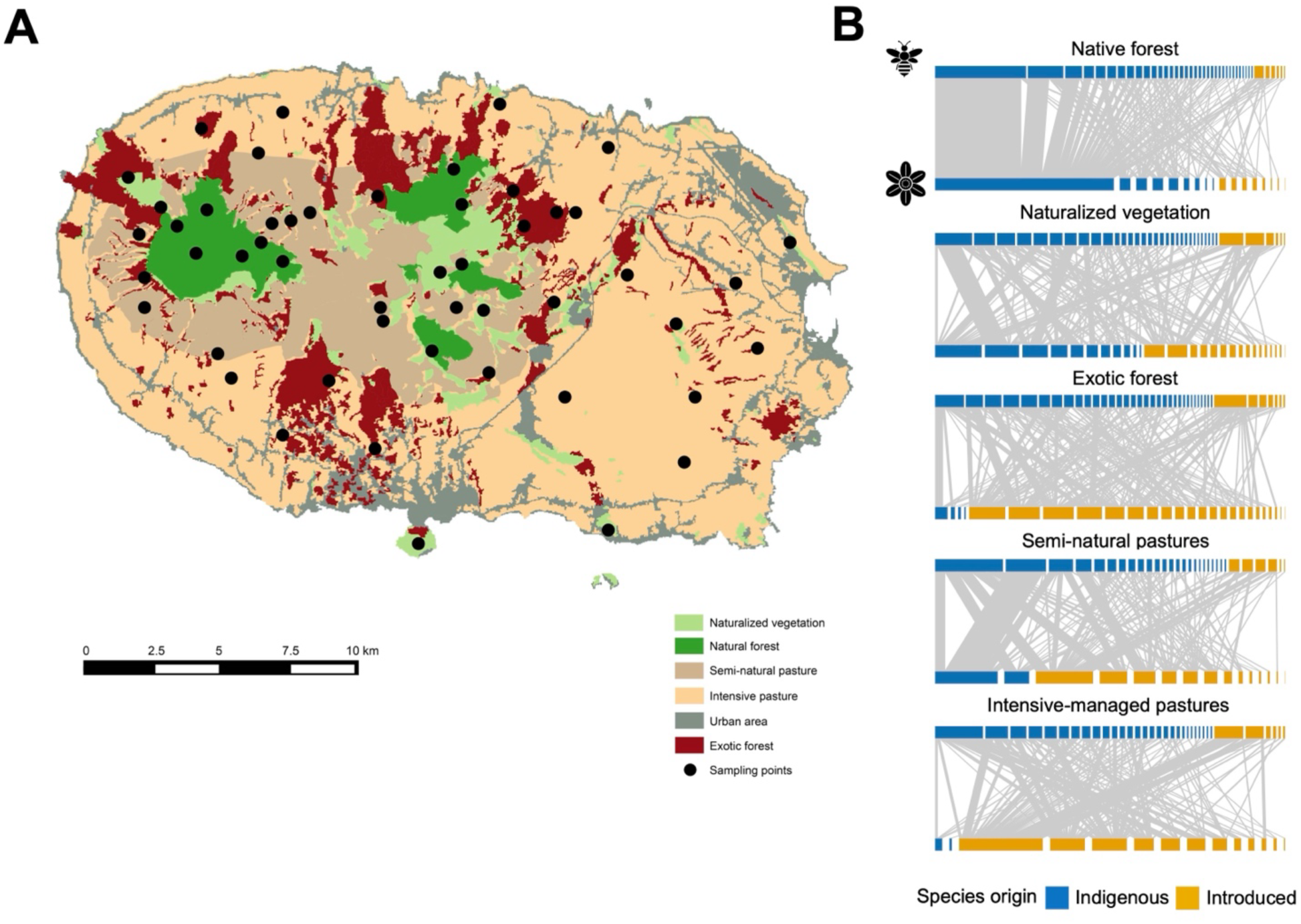
(A) Land use distribution map of Terceira Island with the selected sampling sites (as black dots) covering native forests, naturalized vegetation areas, exotic forests, semi-natural pastures and intensively managed pastures. Cartographic information from DROTRH (2008) and Gaspar (2007), see also Gaspar et al. (2011). (B) Graphical representation of the plant-pollinator networks for the five land-use types.

In the laboratory, collected specimens were sorted first into morphospecies and later identified to species under the supervision of PAVB, following the taxonomic nomenclature in Borges et al. (2010). When species-level identification could not be achieved, individuals were identified to the lowest taxonomic unit possible and classified as morphospecies; this applied to 14% of insect species and 0% of plant species. All species were classified as indigenous or exotic species based on Weissmann et al. (2017) and Borges et al. (2022) for the insects and Borges et al. (2010) for the plants. Importantly, at the time of species identification, uncertainty existed regarding the *Bombus* species present on Terceira, and all specimens were recorded as *Bombus* spp. Following re-evaluation of the specimens and additional sampling (Boieiro et al. 2025a, b), it is likely that most specimens belonged to the introduced species *Bombus terrestris*. In the following text, we retain the denomination *Bombus* spp. and consider all specimens as introduced. Concerning indigenous species, they may be endemic (i.e. found only in the Azores) or native non-endemic (i.e. species that colonized the Azores by natural long-distance dispersal mechanisms and still are found outside the Azores). Exotic species are those whose original distribution range did not include the Azores and which have arrived as a consequence of human activities. In the following analyses, we combined data from the same transects of both years (2013 and 2014) to obtain a better sampling completeness as in Picanço et al. (2017a). We finally built one network for each habitat as the quantitative matrix of interaction between plants (P) and flower-visiting insect (A) species and using the number of visits as link weight (Figure 1). We also built a network at the scale of the island Terceira by pooling the 50 transects together.

### Data analysis

All the following statistical analyses were carried out using the R software version v.4.4.0 (R Core Team, 2024). To assess how thoroughly each network was sampled, we calculated the sample coverage for links (Chiu et al. 2023). Sample coverage for links is a frequency-weighted measure of completeness: it estimates the proportion of interaction events in the network that belong to the links detected in our sample. Rather than indicating how many links remained undetected, coverage reflects how well the interactions are represented in each habitat. Coverage was calculated using the R package (Chiu et al. 2023)

### Quantifying variation in species composition and interactions along the land-use gradient

We identified for each habitat network the number of interactions and visits involving the following combinations: (1) indigenous flower-visiting insects and indigenous plants, (2) indigenous insects and introduced plants, (3) introduced insects and indigenous plants, and (4) introduced insects and introduced plants (Figure 1). To assess how the species and interaction composition of the plant-pollinator networks differ between the different land-use types, we calculated beta diversity between land-use types following the method of Novotny (2009), which describes the processes generating dissimilarities between networks. Network dissimilarity between plant and flower-visiting insects was decomposed into the contributions from species dissimilarity (β_S_) and interaction dissimilarity (β_WN_). Interaction dissimilarity was further partitioned into the effects of species turnover (βST) and interaction rewiring (β_OS_; i.e. species present in both networks but switching interaction partners). The contribution of species turnover to interaction dissimilarity was also decomposed into turnover of insects (β_ST_A), plants (β_ST_P), and both plants and insects simultaneously (β_ST_PA). All metrics were calculated using the *betalinkr* function from the *bipartite* R package (Dormann et al., 2009).

### Quantifying change in network structure along the land-use gradient

To characterize each network structure, we calculated commonly used descriptors that reflect their connectivity, distribution of interactions and specialization: connectance, weighted nestedness, quantitative modularity, network specialization (H^2^’) and specialisation asymmetry. Connectance is the realized proportion of possible interactions, calculated as the number of interactions divided by the size of the network. Quantitative modularity identifies modules in weighted bipartite networks, where each module consists of species that interact more frequently within the module than with species in other modules (Beckett, 2016; Dormann & Strauss, 2014). We conducted quantitative modularity analyses on each network to estimate (a) a measure of modularity (i.e. a value of likelihood) and (b) a number of modules using the quantitative DIRTLPAwb+ algorithm (Beckett, 2016) implemented in the R package *bipartite* (Dormann et al. 2009) with the function *computeModules*. Weighted nestedness was quantified for each network using the metric *WNODF* with the function *nested* in *bipartite* (Almeida-Neto & Ulrich, 2011). This metric extends the classical *NODF* metric (Almeida-Neto et al., 2008) to weighted networks, measuring the extent to which rows and columns display decreasing marginal totals. Network specialisation was calculated using the metric H^2’^, a measure of the degree of niche divergence among species obtained by comparing the observed value with an expected probability distribution of interaction frequencies which assumes that all species interact with their partners in proportion to their observed total frequencies (Blüthgen et al., 2007). It ranges from 0 (low specialisation, high niche overlap) to 1 (high specialisation, low niche overlap). Finally, specialisation asymmetry between plants and flower-visiting insects was calculated as the difference between their standardised specialisation values, with positive values indicating greater specialisation in insects and negative values indicating greater specialisation in plants (Blüthgen et al., 2007).

The significance of modularity and *WNODF* values was assessed against 100 randomizations using a null model. To randomize the networks, we used the *r2dtable* algorithm from the R package *vegan* (Oksanen et al., 2007), which shuffles individual interactions while preserving the total number of interactions per species (i.e., row and column marginal sums). The metrics modularity and *WNODF* values were considered significant if they did not overlap 95% of confidence intervals of the randomized values.

### Quantifying species role along the land-use gradient

To classify each species based on its role in the landscape, we applied the framework developed by Hackett et al. (2019), which builds directly on the approach of Guimerà and Amaral (2005). While the original method defines species roles within and across network modules, Hackett et al. (2019) adapted it to landscape-scale networks, replacing modules with habitat types. This adaptation enables the identification of species roles within a meta-network of habitats. Four broad species roles are distinguished in this framework: (i) habitat hubs are important species due to multiple and/or strong links in a specific habitat, (ii) landscape hubs are important species in multiple habitats, (iii) habitat connectors are found more equally interacting in multiple habitats but are involved in fewer interactions than hubs, and (iv) peripheral species are found predominantly in one habitat and are involved in fewer interactions than hubs.

To classify species, we characterise each species relative importance within habitats with respect to their habitat generalism - i.e. how evenly its interactions are distributed among habitats. This is captured by the relative values of two indices which are: (i) The within-habitat weight *z_i_* of species *i* which measures how important each species is in the landscape, due to strong and/or multiple interactions and it is calculated as follows:

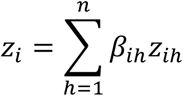

where the coefficient *β_ih_* of species *i* to habitat *h* is the proportion of species *i*’s interactions that occur within habitat *h* and *z_ih_* is the *z*-score of species *i*’s weight - sum of the interaction - in habitat *h*, and it measures how strongly it is connected in the focal habitat and (ii) the among-habitat connectivity *c_i_* which measures how evenly distributed the interactions of species *i* are in the landscape, and is calculated as:

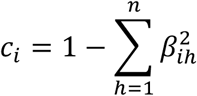

with *c_i_* tending towards 1 if interactions are distributed evenly among the habitats and towards 0 if interactions occur in a single habitat. The four defined species roles can be demarcated in the {*z_i_*, *ci*}-space: peripheral species have a low *z_i_* and a low *c_i_*, habitat connectors have a low *z_i_* and a high *cᵢ*, habitat hubs have a high *zᵢ* and a low *c_i_* and landscape hubs have a high *zᵢ* and a high *cᵢ* (Fig. 2). To evaluate whether species exhibit ‘low’ or ‘high’ values for within-habitat weight *zᵢ* and among-habitat connectivity *cᵢ*, we followed the threshold criteria proposed by Hackett et al. (2019). For within-habitat weight *zᵢ*, which is based on standardized scores, the natural threshold above which *z_i_* is considered significantly higher than the community average is 2.576. This corresponds to a significance level of α = 0.005, thereby controlling for Type I error. For among-habitat connectivity *cᵢ*, species are defined as “connectors” if their highest habitat belonging coefficient *βᵢₕ*is below 60% - that is, less than 60% of their interactions occur within a single habitat type. To determine the corresponding threshold for *cᵢ*, we calculated the maximum *cᵢ* value for species with up to 60% of interactions confined to one habitat. In a meta-network comprising five habitats, this threshold is 0.6, being equal to 60% purely by chance (Hackett et al. 2019). Additionally, we tested whether insects and plants differed in their within-habitat weight (zᵢ) and among-habitat connectivity (cᵢ) using Generalised Linear Model with Gaussian distribution, in which we also included the distinction between native and introduced species as a crossed factor.

**Figure 2.**
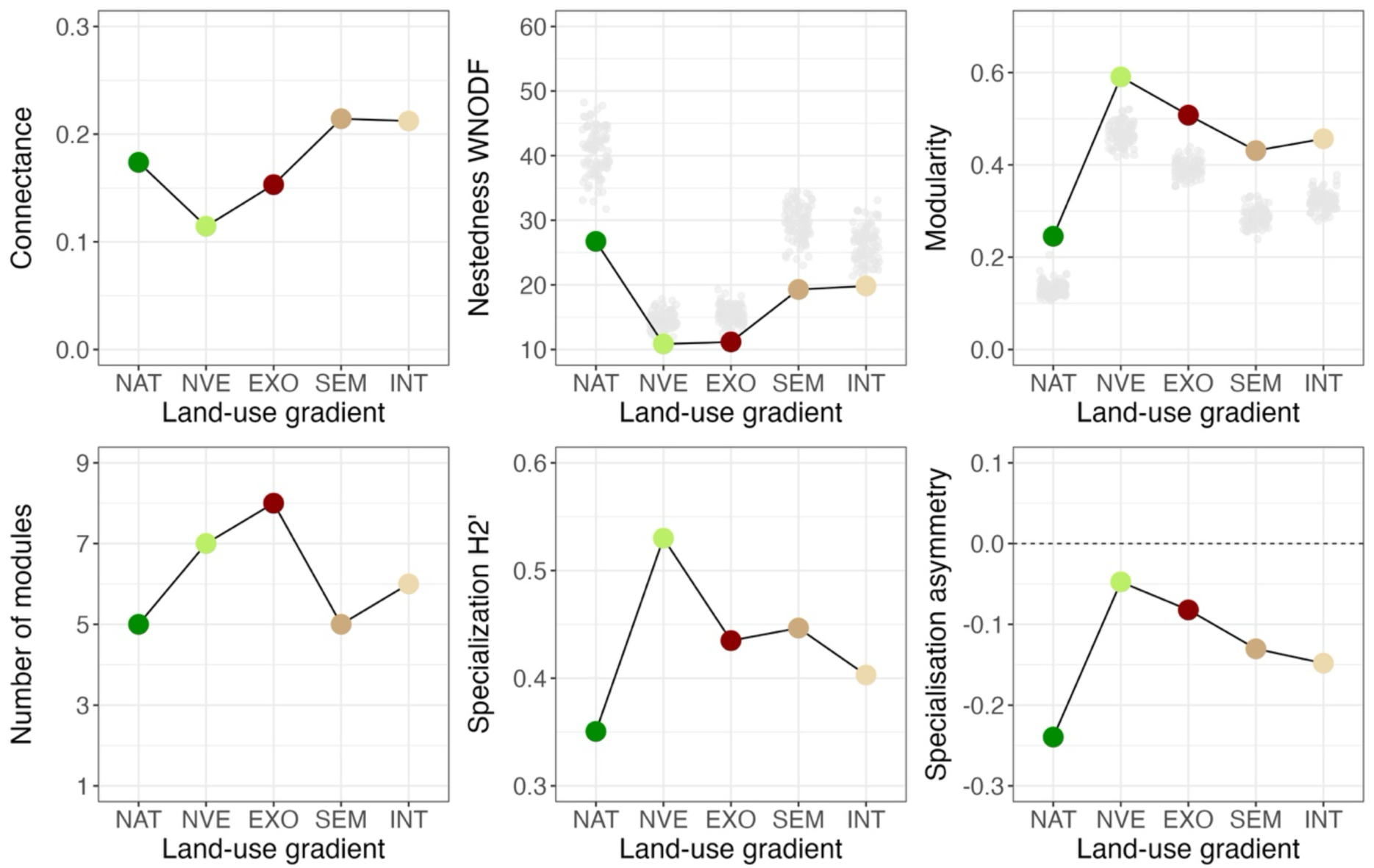
Variation of the network metrics along the land-use gradient. (A) Connectance, (B) Nestedness *WNODF* (C) Quantitative modularity (D) Number of modules (E) Specialisation H^2’^and (F) Specialisation asymmetry. For Nestedness *WNODF* and Quantitative modularity, grey dots indicated the null values generated using the null model *r2dtable* for each land-use type. For nestedness, all observed values were significantly lower than the null expectation while for modularity, all observed values were significantly above the null expectation (Table S2). NAT = Native forest; NVE=Naturalized vegetation; EXO = Exotic forest; SEM = Semi-natural pasture and INT = Intensive-managed pasture.

Finally, we quantified partner similarity across habitats for each insect and plant species, measuring how consistently a species interacts with the same partners across the landscape. Following Hackett et al. (2019), we compared each species’ set of partners within each land-use type to its aggregated set of partners in the meta-network using the Morisita–Horn dissimilarity index. Partner similarity for each species was then calculated as the mean similarity across the five land-use types, where higher values indicate consistent partner associations and lower values indicate partner turnover across habitats. This metric was computed only for species occurring in at least two land-use types. We further tested whether insects and plants differed in their partners’ similarity using Generalised Linear Models (GLM) with Gamma distribution and log-link, in which we also included the distinction between native and introduced species as a crossed factor. We also tested whether, for both insects and plants, species with broader distribution ranges across the land-use gradient exhibit greater resource generalism using Spearman’s correlation between partner similarity and among-habitat connectivity (*c_i_*).

### Sensitivity analyses

As a sensitivity analysis, all analyses detailed above were also performed separately for the 2013 and 2014. In particular, we re-ran the GLMs for within-habitat weight (zᵢ), among-habitat connectivity (cᵢ) and partner fidelity, testing the effects of species level (insects vs. plants) and species origin, and extended these models to also compare the two years. The models therefore included species level, origin and year, as well as all possible interactions among them.

## RESULTS

A total of 54 flower-visiting insects and 48 plant species were recorded, including 8 (15%) and 35 (73%) introduced species, respectively (Table 1). Together, these species accounted for 452 interactions and a total of 2,134 visits. Across the entire gradient, the most frequently observed insect species were the beetle *Anaspis proteus* (352 visits) and the fly *Sepsis neocynipsea* (362), both indigenous. Among plants, the most frequently visited species were the indigenous Asteraceae *Tolpis azorica* (541), endemic to Azores and the introduced Fabaceae *Trifolium repens* (205). The two most frequent interactions were also between indigenous insects with an endemic plant [i.e. *Anaspis proteus* and *Tolpis azorica* (250 visits)] and with an indigenous plant [i.e. *Sepsis neocynipsea* and the indigenous Rosaceae *Potentilla erecta* (64)]. Sample coverage of links was high and similar across all habitats: 93% in native forest, 82% in naturalized vegetation, 80% in exotic forest, 88% in semi-natural pastures and 85% in intensively managed pastures. Because the land-use networks captured a comparable proportion of interactions, differences between land uses are unlikely to reflect differences in sampling completeness.

**Table 1.** Summary of descriptive statistics for the constructed networks. The table includes the abbreviations (Abbr) of land-use types used in the text and figures, the number of species recorded in each network, and the percentage of introduced species (INT) for both plants and flower-visiting insects. For each network, we also report the total number of interactions, the number of visits, and the dominant type of interaction—either between indigenous insects and indigenous plants (IND ↔ IND) or between indigenous insects and introduced plants (IND ↔ INT). Interactions involving introduced insects with either indigenous or introduced plants accounted for only a negligible fraction of the interactions in all networks. The proportions of interactions and visits between pollinators and plants for all combinations of origin are detailed in Table S2.

| Networks (Rank) | Abbr | Plants |  | Flower-visiting insects |  | # interactions (visits) | Dominant interactions |  |
| --- | --- | --- | --- | --- | --- | --- | --- | --- |
|  |  | # species | % INT | # species | % INT | All types | Insects → Plants | % of interactions (% of visits) |
| Native forest (1) | NAT | 17 | 47 | 41 | 15 | 114 (714) | IND → IND | 67 (86) |
| Naturalized vegetation (2) | NVE | 27 | 59 | 37 | 16 | 110 (329) | IND → IND | 53 (56) |
| Exotic forest (3) | EXO | 26 | 85 | 40 | 17 | 147 (388) | IND → INT | 75 (72) |
| Semi-natural pasture (4) | SEM | 15 | 87 | 30 | 20 | 90 (378) | IND → INT | 69 (52) |
| Intensive-managed pasture (5) | INT | 14 | 66 | 34 | 18 | 101 (325) | IND → INT | 83 (75) |

### Variation in species composition and interactions along the land-use gradient

Along the land-use gradient, the number of visited plants recorded ranged from 17 in native forest and 14 in intensively managed pasture to 27 and 26 in naturalized vegetation and exotic forest, respectively—both positioned at mid-gradient intensity (Figure 1B, Table 1). In contrast, the proportion of introduced plants in the networks increased as disturbance intensity increases, from 47% in native forest to 87% and 85% in the two most disturbed land-use types (Table 1). For insects, the trends were less pronounced: the number of flower-visiting species fluctuated between 30 in semi-natural pasture and 41 in native forest, while the proportion of introduced species remained low across the gradient, ranging between 15% and 20% (Table 1). The variation of the proportions of the different interaction types showed marked patterns along the gradient. In less disturbed habitats, most interactions involved indigenous insects and indigenous plants, whereas in more disturbed habitats, interactions were dominated by indigenous insects and introduced plants. A similar pattern was observed when considering the number of visits instead of the number of interactions (Table 1).

However, interactions and visits involving introduced insects were negligible in all land-use types (Table S2).

Our β-diversity analysis between pairs of land-uses revealed that interaction dissimilarity (β_WN_) was the main contributor to network differences over species dissimilarity (β_S_) along the land-use gradient (Table 2). Overall, both β_WN_ and β_S_ tended to increase as land-use types became more distant in disturbance intensity. However, the highest β_S_ and β_WN_ values were observed between exotic forest and semi-natural pastures (β_S_ = 0.46; β_WN_ = 0.92), highlighting a strong shift in species and interaction composition at intermediate gradient intensity. Interaction dissimilarity (β_WN_) between land-use types was mainly driven by species turnover (mean β_ST_ across pairwise values = 0.64 ± 0.15), where changes in interactions between land uses were due to changes in species composition (Table 2). In contrast, species rewiring, representing cases where species shared between two land-use types shifted their interaction partners, contributed less (mean β_OS_ = 0.23 ± 0.12). Interestingly, β_ST_ was largely dominated by plant turnover (mean β_ST_P = 0.48 ± 0.13). This indicates that the emergence of novel interactions with increasing land-use intensity was mostly driven by changes in plant composition (Table 2). Again, β_ST_P reached its maximum (0.66) at intermediate gradient intensity, between exotic forest and semi-natural pastures. In contrast, insect turnover (β_ST_A) contributed very little to overall β_ST_, averaging only 0.05 ± 0.12 (Table 2).

**Table 2.**
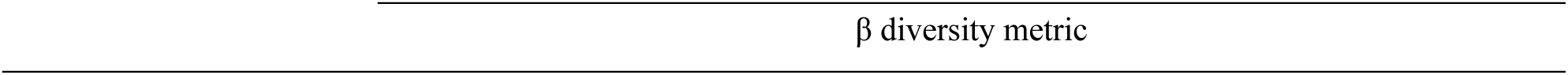
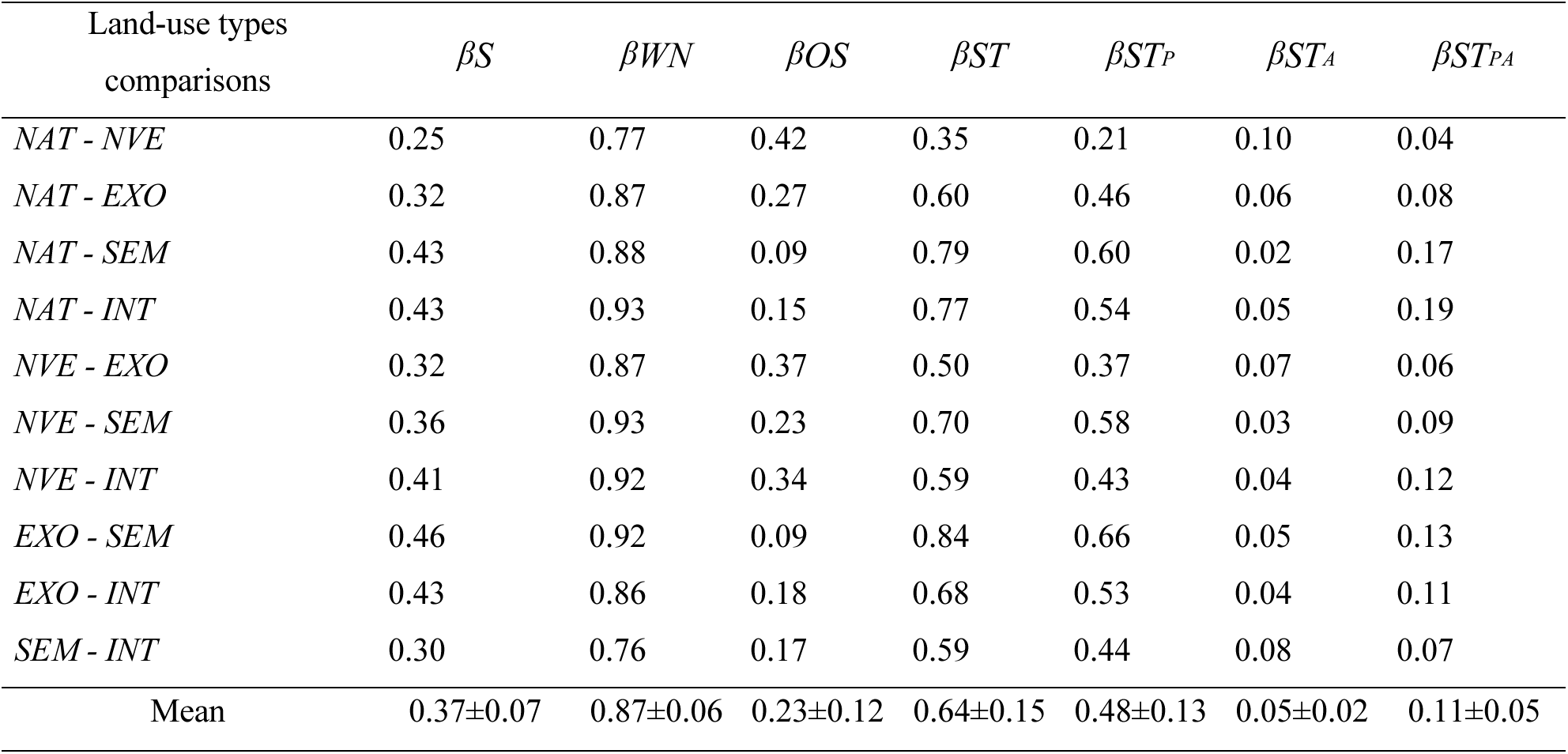
Beta diversity values quantifying change in species and interaction composition between networks of five/different land-use types. Abbreviations: βS, species dissimilarity; β_WN_, interaction network dissimilarity; β_OS_, interaction dissimilarity explained by rewiring among shared species; β_ST_, interaction dissimilarity due to species turnover; β_ST_A, interaction dissimilarity due to insect turnover; β_ST_P, interaction dissimilarity due to plant turnover; β_ST_PA, interaction dissimilarity due to insect and plant turnover. NAT = Native forest; NVE=Naturalized vegetation; EXO = Exotic forest; SEM = Semi-natural pasture and INT = Intensive-managed pasture.

| Land-use types<br>comparisons | $\beta_S$ | $\beta_{WN}$ | $\beta_{OS}$ | $\beta_{ST}$ | $\beta_{ST_P}$ | $\beta_{ST_A}$ | $\beta_{ST_{PA}}$ |
| --- | --- | --- | --- | --- | --- | --- | --- |
| *NAT - NVE* | 0.25 | 0.77 | 0.42 | 0.35 | 0.21 | 0.10 | 0.04 |
| *NAT - EXO* | 0.32 | 0.87 | 0.27 | 0.60 | 0.46 | 0.06 | 0.08 |
| *NAT - SEM* | 0.43 | 0.88 | 0.09 | 0.79 | 0.60 | 0.02 | 0.17 |
| *NAT - INT* | 0.43 | 0.93 | 0.15 | 0.77 | 0.54 | 0.05 | 0.19 |
| *NVE - EXO* | 0.32 | 0.87 | 0.37 | 0.50 | 0.37 | 0.07 | 0.06 |
| *NVE - SEM* | 0.36 | 0.93 | 0.23 | 0.70 | 0.58 | 0.03 | 0.09 |
| *NVE - INT* | 0.41 | 0.92 | 0.34 | 0.59 | 0.43 | 0.04 | 0.12 |
| *EXO - SEM* | 0.46 | 0.92 | 0.09 | 0.84 | 0.66 | 0.05 | 0.13 |
| *EXO - INT* | 0.43 | 0.86 | 0.18 | 0.68 | 0.53 | 0.04 | 0.11 |
| *SEM - INT* | 0.30 | 0.76 | 0.17 | 0.59 | 0.44 | 0.08 | 0.07 |
| Mean | 0.37±0.07 | 0.87±0.06 | 0.23±0.12 | 0.64±0.15 | 0.48±0.13 | 0.05±0.02 | 0.11±0.05 |

### Network structure along the land-use gradient

Our results show that land uses at intermediate disturbance intensities consistently exhibit distinct characteristics compared with both the least and most disturbed land uses. Connectance and nestedness were lowest in naturalized vegetation and exotic forest (Figure 2A, B), while modularity and the number of modules peaked in these same land-uses with a maximum of 8 modules in both land-uses (Figure 2C, D). Across all land-use types, networks were significantly anti-nested with observed nestedness lower than expected by chance and more modular than expected by chance (All land-uses: P = 0.01, Table S3). Regarding specialization, the native forest exhibited the lowest H2’ value (H2’ = 0.35), indicating the least specialized network of the gradient, whereas the highest value was observed in naturalized vegetation (H2’ = 0.53) (Figure 2E). Specialization asymmetry was negative across all land-use types, suggesting that plant species were generally more specialized than flower-visiting insects (Figure 2F). This asymmetry was most pronounced in native forest, while naturalized vegetation exhibited the least asymmetry among land uses.

### Species roles along the land-use gradient

The evaluation of the structural species role in the {*z_i_, c_i_*} space, defined by within-habitat weight (*zi*) and among-habitat connectivity (*c_i_*) (Figure 3A) showed that, among insects 52% of the species (28 species) were classified as peripheral, 46% (25 species) as habitat connectors, and only 2% (one species) as a marginal habitat hub (Figure 3B). Among plants, the majority were peripheral (81%; 39 species), with only 10% (5 species) identified as habitat connectors and 8% (4 species) as habitat hubs (Figure 3C). No insect or plant species was identified as a landscape hub. Analyses of each dimension of the {*z_i_, c_i_*} space confirmed the above pattern: pollinator insects had lower within-habitat weight (*zi*) than plant species whereas insects exhibited higher among-habitat connectivity (*c_i_*) than plants (Figure 4A, B, Table S4). No significant differences were found between indigenous and introduced species as well as no interaction between species levels and origin either within-habitat weight or among-habitat connectivity for both insects and plants (Table S4).

**Figure 3.**
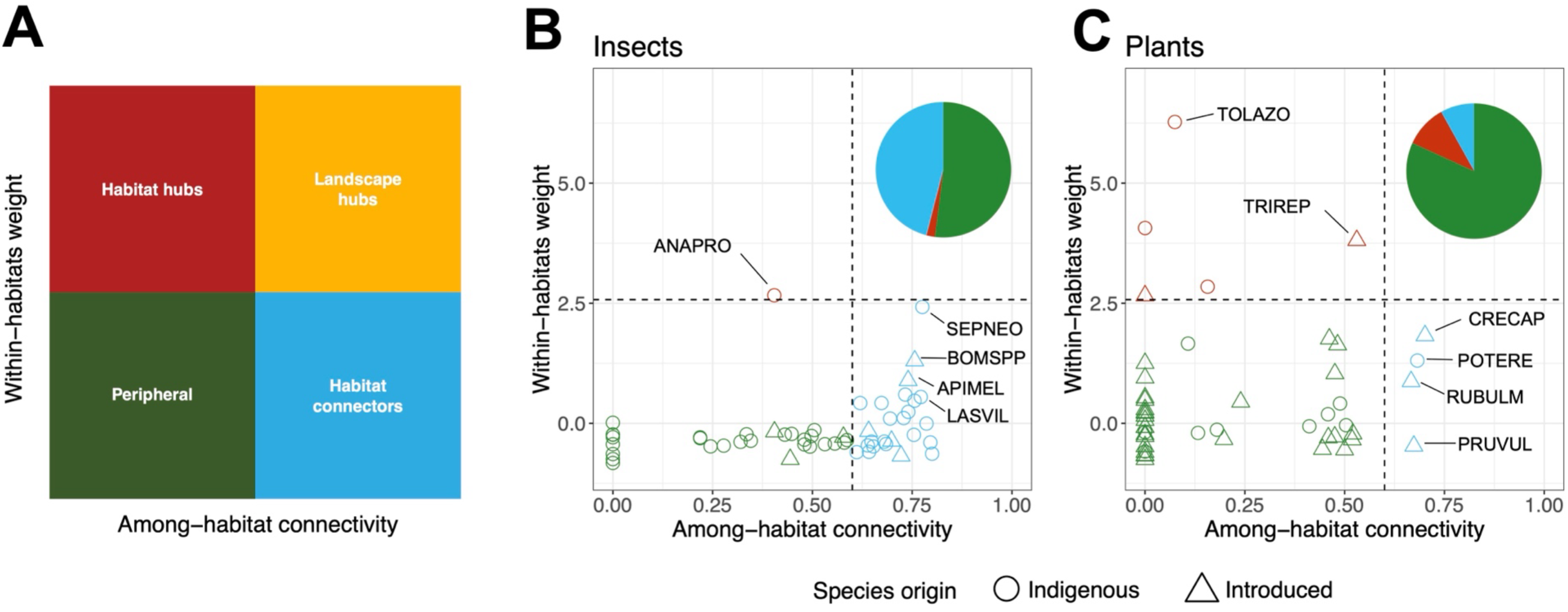
(A) Distribution of species roles in the landscape matrix according to their coordinates in the {zi, ci}-space (within-habitat weight against among-habitat connectivity). (B, C) Species distribution in the {zi, ci}-space according to the quantitative networks for (B) flower-visiting insects and (C) Plants. The top right pie charts indicate the proportion of species in each role defined in (A). Symbols distinguish indigenous (open circles) and introduced species (upward triangles). Symbol colour refers to the assigned landscape role in (A). Flower-visiting Insects: ANAPRO = *Anaspis proteus* (Coleoptera); SEPNEO= *Sepsis neocynipsea* (Diptera); BOMSPP= *Bombus* spp. (Hymenoptera); APIMEL= *Apis mellifera* (Hymenoptera); LASVIL = *Lasioglossum villosulum* (Hymenoptera). Plants: TOLAZO = *Tolpis azorica* (Asterales); TRIREP = *Trifolium repens* (Fabales); CRECAP = *Crepis capillaris* (Asterales); POTERE = *Potentilla erecta* (Rosales); RUBULM = *Rubus ulmifolius* (Rosales); PRUVUL = *Prunella vulgaris* (Lamiales).

**Figure 4.**
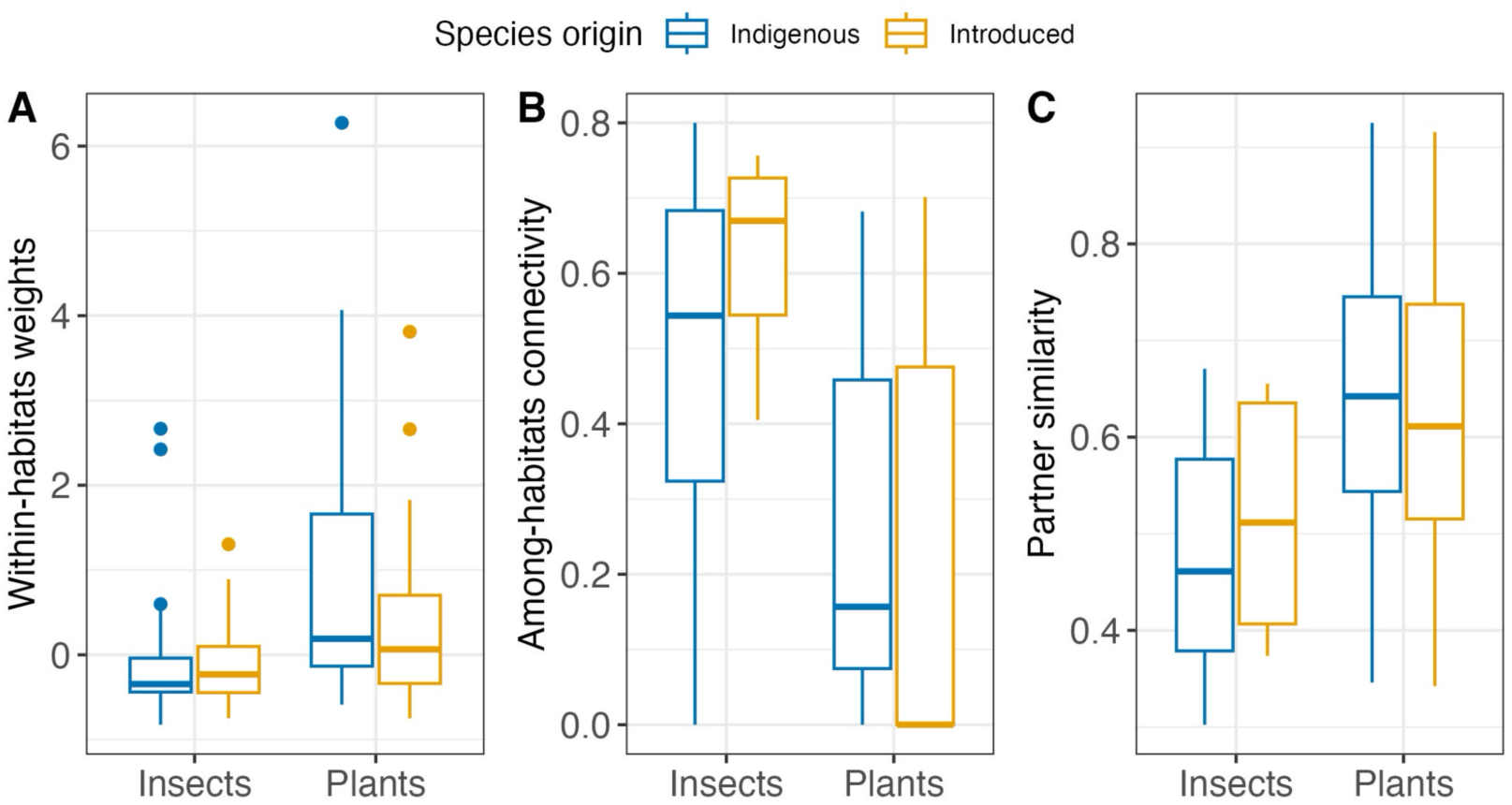
Variation in (A) within-habitat weight (*z_i_*), (B) among-habitat connectivity (*c_i_*), and (C) partner similarity between flower-visiting insects and plants. For each group, differences between indigenous and introduced species are also shown. In all cases, insects and plants differed significantly (Table S3), whereas no significant differences were detected between indigenous and introduced species within each group and for each variable.

Analysis of the distribution of visits across land-use types by the main habitat connectors among pollinator insects (Figure 5) confirmed that these species use a broad range of habitats, visiting plants in all five land-use types. Moreover, the origins of the plants they visited indicate that these connector species can shift from indigenous to introduced plant partners along the land-use gradient.

**Figure 5.**
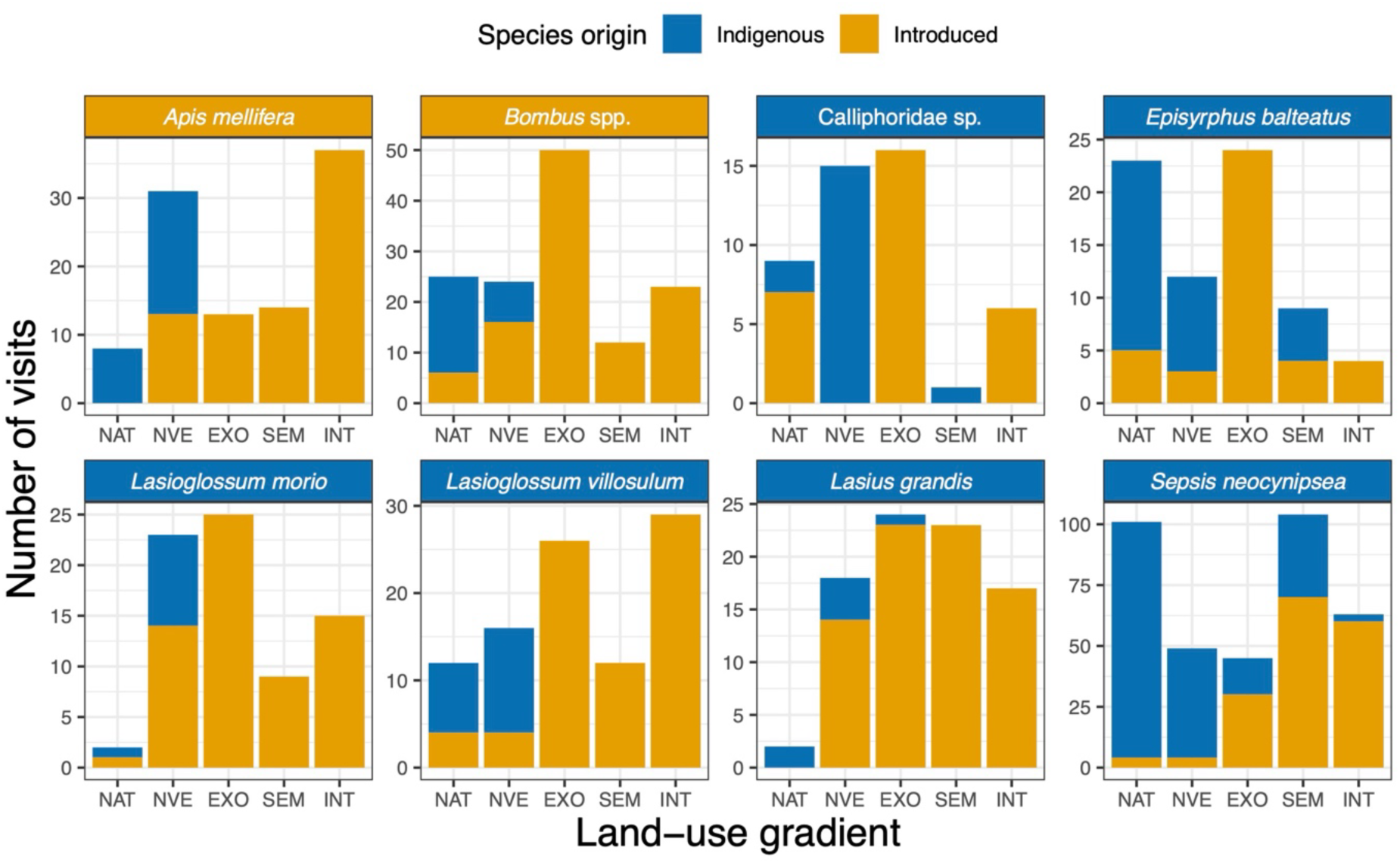
Examples of the eight most important habitat connectors among flower-visiting insects based on their position in the {*z_i_, c_i_*} space. For each insect species, the distribution of visits across land-use types is shown, illustrating their broad habitat range and their ability to shift from indigenous to introduced plant partners along the gradient. NAT = Native forest; NVE=Naturalized vegetation; EXO = Exotic forest; SEM = Semi-natural pasture and INT = Intensive-managed pasture.

Partners’ similarity across land uses, measured as the average similarity in partner composition across land uses, was calculated only for species occurring in at least two land-use types: 46 insect species (85%) and 25 plant species (52%), indicating that half of the plant species seem to be restricted to a single land use, compared with only 15% of the insect species. Overall, partner similarity was lower for insects than for plants (Figure 4). Again, we found no significant differences between indigenous and introduced species in partner similarity as well as no effect of the interaction between species levels and species origin (Table S4). Partner similarity for insects was significantly and negatively correlated with among-habitat connectivity (ci) (Spearman correlation: r=-0.43; P = 0.002), suggesting that species connecting habitats use different sets of resources in each land-use (i.e. diet generalist). In contrast, no correlation was reported for plants (r= 0.36; P = 0.08) (Figure 6).

**Figure 6.**
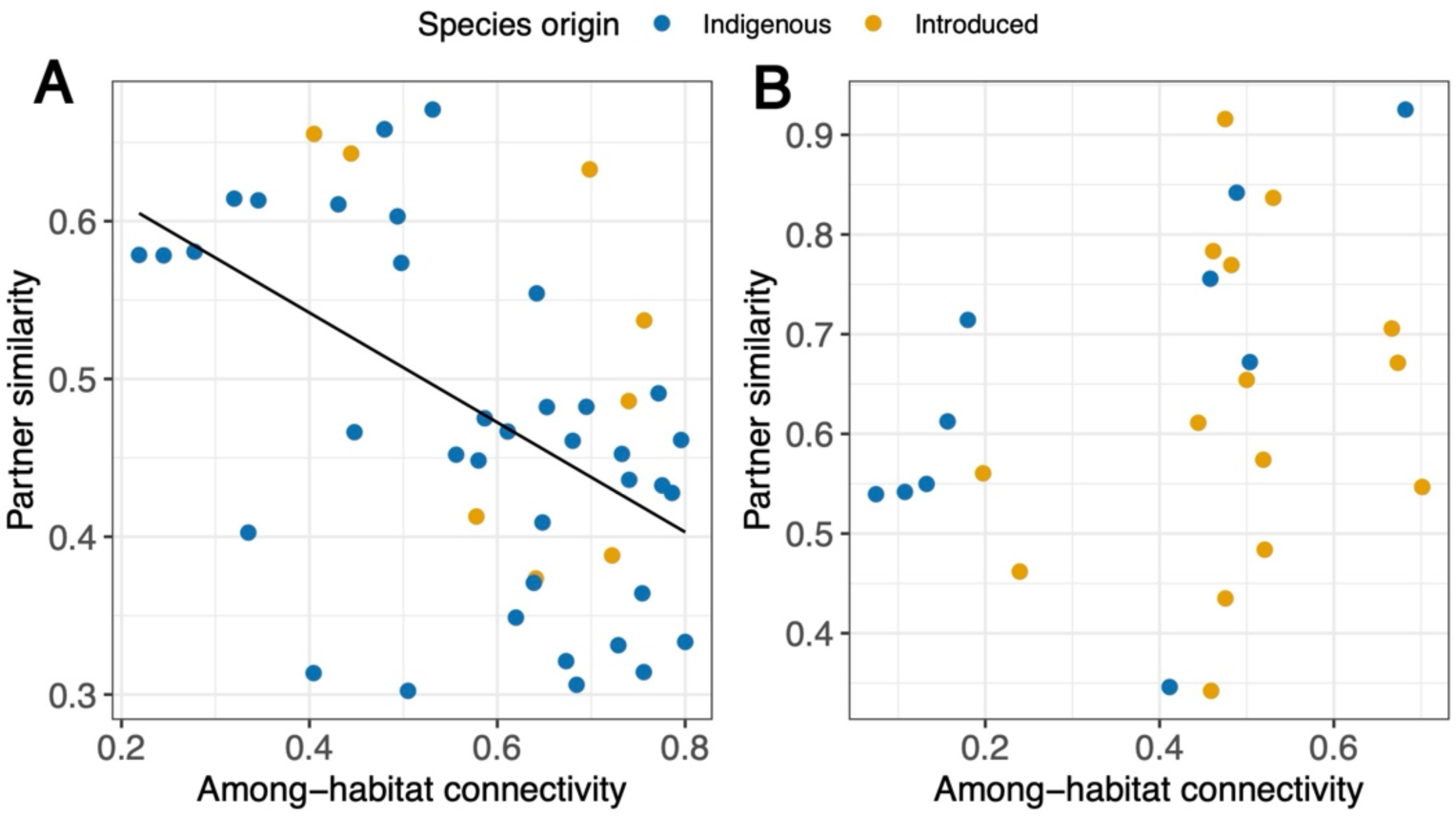
Relationship between partner similarity and among-habitat connectivity for (A) flower-visiting insects and (B) plants. For both groups, the data points for indigenous and introduced species are indicated with different colours. The regression line for (A) indicated a significant negative relationship according to the Spearman correlation.

### Sensitivity analyses

Overall, when the two years were analysed separately, they did not show any specific differences between them (Tables S2-3, S5-S8; Figures S1-S7) or with the combined dataset (results detailed above). More specifically, when we tested the effects of species level (insect vs plants), origin (indigenous vs introduced), year (2013 vs 2014) and all possible interactions between them on among-habitat connectivity (cᵢ), within-habitat weight (zᵢ) and partner similarity (Table S5), species level was the only significant effect, except for within-habitat weight for which the effect of origin was marginal significant (Table S5).

## DISCUSSION

Our study demonstrates that land-use change on Terceira has led to a strong reorganization of plant–pollinator networks, but not to a collapse: species and interaction diversity, as well as overall network topology, remain largely unchanged between pristine and highly disturbed habitats. Instead, intermediate habitats, such as naturalized vegetation and exotic forests, emerged as areas of network reassembly. Changes along the gradient were driven primarily by plant turnover, while flower-visiting species, mostly indigenous, persisted across habitats by shifting their interaction partners. These pollinator species can be seen as super-generalists that act as interaction bridges across habitats, allowing continuity of pollination function across the landscape despite changes in land-use.

This study complements the growing body of research on plant-pollinator networks in the Azores and further highlights this archipelago as a valuable model system for studying pollination networks on oceanic islands under human disturbance (Boieiro et al., 2025a, 2025b, 2025c; Olesen et al., 2002; Picanço et al., 2017a, 2017b; Weissmann & Schaefer, 2017). It also provides the first comprehensive assessment of how plant-pollinator network structure reorganizes along a full land-use gradient on an oceanic island, integrating changes in network topology with shifts in species roles across habitats.

### Interaction turnover along the gradient is driven by changes in plant species composition

The establishment of the land-use gradient on Terceira Island resulted in a strong interaction dissimilarity (β_WN_) between habitats, reaching 87% on average. As predicted, this reshuffling of interactions was mostly driven by plant turnover (β_ST_P), with only a small contribution from insect turnover (β_ST_A). Plant turnover across habitats documented herein is consistent with previous studies in the Azores (e.g. Elias et al. 2016; Leo et al. 2024). The rewiring component (β_OS_, quantifying the switch of interaction partners for species present in both networks of the pair) contributed little to interaction dissimilarity. This minor role of rewiring has already been reported along gradients of elevation, urbanisation, and landscape simplification (Maurer et al., 2024; Simanonok & Burkle, 2014; Sponsler et al., 2022; White et al., 2022, but See CaraDonna et al., 2017). One explanation for this pattern is that environmental filtering acting along ecological gradients primarily changes the species composition of at least one of the two network levels (mostly plants; e.g. Maurer et al., 2024; Sponsler et al., 2022). Change in interaction composition therefore results primarily from the loss or gain of species, and not from the rewiring of interactions among species co-occurring across the gradient (Simanonok & Burkle, 2014; Tylianakis & Morris, 2017).

Importantly, plant turnover was mostly due to the establishment of introduced plants into the networks of human-made habitats, being the dominant plant species in the more disturbed habitats of the Azorean Islands (Silva & Smith, 2004). In contrast, plant turnover was accompanied by pollinator persistence across land-uses (β_ST_A = 5%) with a dominance of indigenous pollinator insects. This persistence across contrasting habitats, together with their broad use of locally available floral resources, is consistent with ecological release, whereby indigenous island pollinators expand their realised niches and adopt highly generalised foraging strategies (Picanço et al., 2017a). Indeed, in all habitats, the majority of the interactions and visits involved an indigenous insect (Table 1), visiting either an indigenous plant in the two least disturbed habitats (native forest and naturalized vegetation) or an introduced plant in the more disturbed habitats (i.e., exotic forest, semi-natural pasture, and intensively managed pasture). Introduced pollinator species remained marginal throughout the gradient (Picanço et al., 2017a), although both *Apis mellifera* and *Bombus* spp. were found to be substantial contributors to plant visitation in most habitats (Figure 5), as reported by Boieiro et al. (2025a).

### Network structure responds non-monotonically to land use

Network structure varied non-monotonically along the gradient (see also Tylianakis & Morris, 2017). Connectance and weighted nestedness were lowest, and modularity and the number of modules highest, in the intermediate habitats (naturalized vegetation and exotic forest) rather than at the least or most disturbed ends of the gradient. These mid-gradient networks were structurally distinct rather than simply degraded versions of native forest: they hosted the highest number of plant species, mixing plants from both indigenous and introduced origin, and were also among the richest for pollinators, leading to a high number of distinct interactions (Table 1). These features were also associated with the highest modularity values recovered along the gradient (Figure 2C). When floras of different origin, trait composition, or phenology coexist, their associated pollinator interactions are not necessarily randomly distributed across the network; they can instead cluster into distinct modules. For instance, indigenous generalist insects, which could potentially interact with both native and introduced plants, must show some degree of interaction selectivity toward one pool over the other (Timoteo et al. 2018). We suggest that introduced plants, which often present novel floral traits or phenology relative to indigenous plants (Lázaro-Lobo et al., 2025), may recruit certain indigenous insects, reducing their visitation to native plants through closer trait/phenological matching with the introduced flora (Albrecht et al., 2014; Kandori et al., 2009; Lopezaraiza–Mikel et al., 2007, Timoteo et al. 2018). The result is fewer cross-pool interactions and denser within-pool interactions, reinforcing module boundaries. This interpretation is supported by complementary specialisation metrics: high modularity, especially in naturalized vegetation, was accompanied by high network specialisation and low specialisation asymmetry (Figure 2E-F). This indicates that both plants and insects are, on average, specialised enough within the habitat (i.e. restricting their interactions to a subset of potential partners) to generate distinct modules.

Surprisingly, all networks were found to display nestedness lower than expected by chance and modularity higher than expected by chance. This might indicate that, on Terceira Island, disturbance did not drive the system toward the generalized, nested structures typically expected for mutualistic and island pollination networks (Bascompte et al., 2003; Olesen & Jordano, 2002; Trøjelsgaard & Olesen, 2016). This result is also consistent with the evidence that nestedness in mutualistic networks may be weaker than long assumed (Staniczenko et al., 2013).

### Indigenous super-generalist pollinators as landscape connectors

The analysis of species structural roles (Hackett et al., 2019) enabled us to precisely identify the role of pollinator insects in connecting networks across the land-use gradient. We found that nearly half of the pollinator insects were habitat connectors, distributing their interactions across most habitats, whereas most plants were restricted to a few habitats, often occurring in a single one; insects showed higher among-habitat connectivity (c_i_) and lower within-habitat weight (z_i_) than plants. No insect species qualified as a landscape hub, so network cohesion was maintained mostly by a set of broadly foraging indigenous insects (Figures 3 and 4).

We also demonstrated that these flower-visiting habitat connectors were not tracking a specific resource throughout the landscape. Instead, we found that partner similarity between habitats declined with among-habitat connectivity (c_i_), so the most widely distributed insect species visited different sets of plants in each habitat, behaving as context-dependent generalists interacting with flowers locally available (Poisot et al., 2015). This pattern is consistent with ecological release and density compensation phenomena already documented on species-poor islands (Kaiser-Bunbury et al., 2009; Olesen et al., 2002).

Introduced plant species were integrated into these networks in a similar way to indigenous ones. Indeed, the two groups did not differ in within-habitat weight (z_i_), among-habitat connectivity (c_i_), or partner similarity so, once established, introduced plants were incorporated into the interaction network by indigenous super-generalists in the same way as indigenous plants (Bartomeus et al., 2008; Vilà et al., 2009). This integration has one important consequence: the insects that maintain the pollination of indigenous plants persisting outside their native habitat (Ghisbain et al., 2021) may also facilitate the reproduction and spread of introduced plants (Traveset & Richardson, 2014) in the landscape. Indigenous super-generalists therefore play a central role in these novel networks, contributing both to network resilience and to the integration of introduced species, with a net effect on the ecosystem that remains to be assessed. For instance, the integration of introduced plants into the network can redirect pollinator visitation and change native plant reproduction even when overall interaction networks remain connected (Brett et al., 2024).

### Implications for conservation and restoration on oceanic islands

Our results have several implications for the conservation and restoration of plant-pollinator networks on oceanic islands. First, they highlight the value of conserving indigenous pollinator assemblages even in disturbed habitats (M’Gonigle et al., 2015), since it is the persistence and the foraging flexibility of these species that maintains pollination connectivity across the landscape (Kaiser-Bunbury et al., 2017; Rojas-Sandoval et al., 2026). Second, because interaction turnover is governed by changes in plant composition, conservation programs aiming at restoring native interactions will be most effective if they target the plant community, re-establishing indigenous flora rather than assuming that pollinators alone will rebuild lost links (Menz et al., 2011). Third, our results highlight the importance of giving particular attention to intermediate habitats. Rather than being simple transition zones, naturalized vegetation and exotic forests have structurally distinct networks where indigenous and introduced species coexist and where network reassembly seems to be most intense (Gutiérrez-Chacón et al., 2020; Vasiliev & Greenwood, 2023). Overlooking these habitats would mean disregarding a substantial part of the landscape interaction diversity.

### Limitations

Nevertheless, our study has several limitations. Timed focal observations may disproportionately record common and frequently visiting insects, while rare or temporally restricted interactions may remain undetected (Gibson et al., 2011; Jordano, 2016). Moreover, because visitors were not individually marked, successive observations may include repeated visits by the same forager, potentially inflating the apparent frequency of particular plant–insect links during flower-constant foraging bouts (Dupont et al., 2011). Our conclusions rely on a single island and two field seasons, so the generality of these patterns across the entire archipelago, and between years, remains to be tested. Finally, what we consider as interactions describe only visitation, not effective pollination (King et al., 2013). For instance, no data exist for the Azores on whether recorded visits translate into successful pollen transfer, particularly for introduced plants.

## CONCLUSIONS

Land-use change on Terceira Island reorganizes plant–pollinator networks mainly through plant turnover, whereas pollinator insect assemblages remain comparatively stable across habitats. The replacement of indigenous by introduced plants as land-use intensity increases drives an almost complete turnover of interactions, yet a persistent fauna of indigenous super-generalist pollinators continually rewires its partners by tracking local floral resources, thus maintaining pollination connectivity across the gradient. On species-poor oceanic islands, such broadly foraging indigenous species may constitute an important stabilizing component of the pollination systems, sustaining the continuity of mutualistic function through periods of rapid anthropogenic change.

## AUTHOR CONTRIBUTIONS

François Rigal, Paulo A. V. Borges and Ana Picanço conceived the ideas and designed methodology; Ana Picanço collected the data; François Rigal and Ana Picanço analysed the data with the advices of Sérgio Timóteo and Mário Boieiro; François Rigal led the writing of the manuscript. All authors contributed critically to the drafts and gave final approval for publication.

## ACKNOWLEDGEMENTS

We are grateful to Virgílio Vieira and Matthias Jentzsch who assisted in the identification of the lepidopteran morphospecies and *Xanthandrus azorensis*, respectively. During the preparation of this work the authors use WRITFULL in order to improve the grammar and the readability of the work. After using this tool, the authors reviewed and edited the content as needed, and take full responsibility for the content of the publication.

## DATA AVAILABILITY STATEMENT

Data used in this manuscript are available from the the link: https://doi.org/10.6084/m9.figshare.34572102

## SUPPORTING INFORMATION

Additional supporting information can be found online in the Supporting Information section at the end of this article.

